# A post-decisional neural marker of confidence predicts information-seeking in decision-making

**DOI:** 10.1101/433276

**Authors:** Kobe Desender, Peter Murphy, Annika Boldt, Tom Verguts, Nick Yeung

**Affiliations:** Department of Neurophysiology and Pathophysiology, University Medical Center Hamburg-Eppendorf, Germany; Department of Experimental Psychology, Ghent University, Belgium; Institute of Cognitive Neuroscience, University College London, United Kingdom; Department of Experimental Psychology, University of Oxford, United Kingdom

**Keywords:** confidence, metacognition, decision making, information sampling, information-seeking, error positivity, Pe, EEG, P3

## Abstract

Theoretical work predicts that decisions made with low confidence should lead to increased information-seeking. This is an adaptive strategy because it can increase the quality of a decision, and previous behavioral work has shown that decision-makers engage in such confidence-driven information seeking. The present study aimed to characterize the neural markers that mediate the relationship between confidence and information-seeking. A paradigm was used in which human participants made an initial perceptual decision, and then decided whether or not they wanted to sample more evidence before committing to a final decision and confidence judgment. Pre-decisional and post-decisional ERP components were similarly modulated by the level of confidence and by information-seeking choices. Time-resolved multivariate decoding of scalp EEG signals first revealed that information-seeking choices could be decoded from the time of the initial decision to the time of the subsequent information-seeking choice (within-condition decoding). No above-chance decoding was visible in the pre-response time window. Crucially, a classifier trained to decode high versus low confidence predicted information-seeking choices after the initial perceptual decision (across-condition decoding). This time window corresponds to that of a post-decisional neural marker of confidence. Collectively, our findings demonstrate for the first time that neural indices of confidence are functionally involved in information-seeking decisions.

## Introduction

Humans seek information adaptively to improve the quality of their decisions, for example by requesting expert advice when they lack the relevant domain knowledge or discriminating evidence (for review, see Bonaccio & Dalal, 2006). This tendency has been documented for everyday financial (Yoong & Hung, 2010) and medical (Butler, Danby, Emmison, & Thorpe, 2009) decisions, as well as in more carefully controlled lab settings (Sniezek & Van Swol, 2001). Recent evidence suggests that information seeking depends crucially on explicit representation of decision confidence (Desender, Boldt, & Yeung, 2018). Theoretically, decision confidence has been treated as an internal evaluation signal that can be used to adapt behavior in the absence of external feedback (Meyniel, Sigman, & Mainen, 2015; Yeung & Summerfield, 2012): When confidence in a decision is low, this implies that the probability of a decision being correct is also low, and seeking additional information before committing to a decision might be particularly beneficial. In a previous behavioral study, participants engaged more in information-seeking in conditions associated with low compared to high confidence, despite equal accuracy in both (Desender et al., 2018).

At present, however, it remains unclear how the neural coding of confidence informs the decision to engage in additional information seeking. It has been argued that decision confidence reflects the strength of the evidence in favor of a decision (Vickers, 1979; Zylberberg, Barttfeld, & Sigman, 2012), stressing the importance of pre-decisional evidence in the computation of confidence (Kiani & Shadlen, 2009). On such a view, neural signals involved in the decision process itself should be predictive of information seeking (e.g., the P3 component of the scalp-recorded EEG; O’Connell, Dockree, & Kelly, 2012; Twomey, Murphy, Kelly, & O’Connell, 2015). Alternatively, decision confidence has been quantified as a function of continued evidence accumulation following a decision (Moran, Teodorescu, & Usher, 2015b; Pleskac & Busemeyer, 2010), stressing the importance of post-decisional signals in the computation of confidence. Recently, a post-decisional centro-parietal positivity was found in scalp EEG recordings that reflects post-decisional neural evidence accumulation informing judgments about the accuracy of the preceding decision (Murphy, Robertson, Harty, & O’Connell, 2015). This signal is commonly referred to as the error positivity (Pe; Nieuwenhuis, Ridderinkhof, Blom, Band, & Kok, 2001), and has been shown to reflect fine-grained variations in decision confidence (Boldt & Yeung, 2015).

The current study aimed to identify neural signatures of confidence that are predictive of information seeking behavior. In our experimental paradigm, on each trial participants made an initial perceptual decision about the mean color of eight visual elements, and then decided whether or not to sample additional evidence (at a small cost) before committing to a final decision. Electrophysiological recordings allowed us to evaluate which neural signatures of confidence were related to information-seeking choices. To do so, we relied on time-resolved decoding of EEG data to test for shared neural coding of confidence and decisions to seek additional information. Specifically, we tested whether a multivariate classifier trained to decode confidence from EEG data would be predictive of information-seeking choices. Such between-condition generalization isolates neural processes integral to translating decision confidence into the overt decision to sample additional information. Our main question of interest was whether information-seeking choices could be predicted based on neural markers of confidence observed pre-decisional (P3), post-decisional (Pe), or both.

## Method

### Participants

Seventeen participants (nine males, mean age: 24.1 years, SD = 3.0, range 21 – 32) took part at Oxford University for monetary compensation (£20 plus up to £4.92 dependent on performance, range of the rounded actual payments: £22 − £24). The data of two participants were excluded because accuracy of the primary responses was at chance level (48.9% and 49.6% correct), thus the final sample comprised fifteen participants. All provided written informed consent, reported normal or corrected-to-normal vision and were naive with respect to the hypothesis. All procedures were approved by the local ethics committee.

### Stimuli and apparatus

Stimuli were presented on a gray background on a 20-inch CRT monitor with a 75 Hz refresh rate, using the MATLAB toolbox Psychtoolbox3. Each stimulus consisted of eight colored shapes spaced regularly around a fixation point (radius 2.8° visual arc). To manipulate task-difficulty, the mean and the variance of the eight elements varied across trials. The mean color of the eight shapes was determined by the variable *C*; the variance across the eight shapes by the variable *V*. The mean color of the stimuli varied between red ([1, 0, 0]) and blue ([0, 0, 1]) along a linear path in RGB space ([*C*,0, 1 − *C*]). At the start of the experiment, *C* could take four different values: 0.450, 0.474, 0.526 and 0.550 (from blue to red, with 0.5 being the category boundary), and *V* could take two different values: 0.0333 and 0.1000 (low and high variance, respectively). On every trial, the color of each individual element was pseudo-randomly selected with the constraint that the mean and variance of the eight elements closely matched the mean of *C* and its variance *V*, respectively. Each combination of *C* and *V* values occurred equally often. The individual elements did not vary in shape. Responses were made using a USB mouse and a standard QWERTY keyboard.

### Procedure

Figure 1 shows an example trial during the main part of the experiment. After a 200 ms fixation interval, the stimulus was flashed for 200ms. Participants were instructed to respond as quickly as possible, deciding whether the average color of the eight elements was more blue or more red, by clicking one of two mouse buttons. The mapping between color and response was counterbalanced between participants. After a 200 ms post-response interval there was a choice phase consisting of two conditions: free-choice or no-choice. On free-choice trials (75% of trials), the letters R and S appeared, indicating that participants could either choose to request additional evidence by seeing the stimulus again in an easier version (S), or to give their response (R). They indicated their choice by moving a grey slider up or down with their mouse towards their choice, and confirmed by pressing the spacebar (locations of R and S were fixed across trials and counterbalanced across participants). On no-choice trials (25% of the trials), only an R appeared, and participants were forced to select the option to give their response.

**Figure 1.**
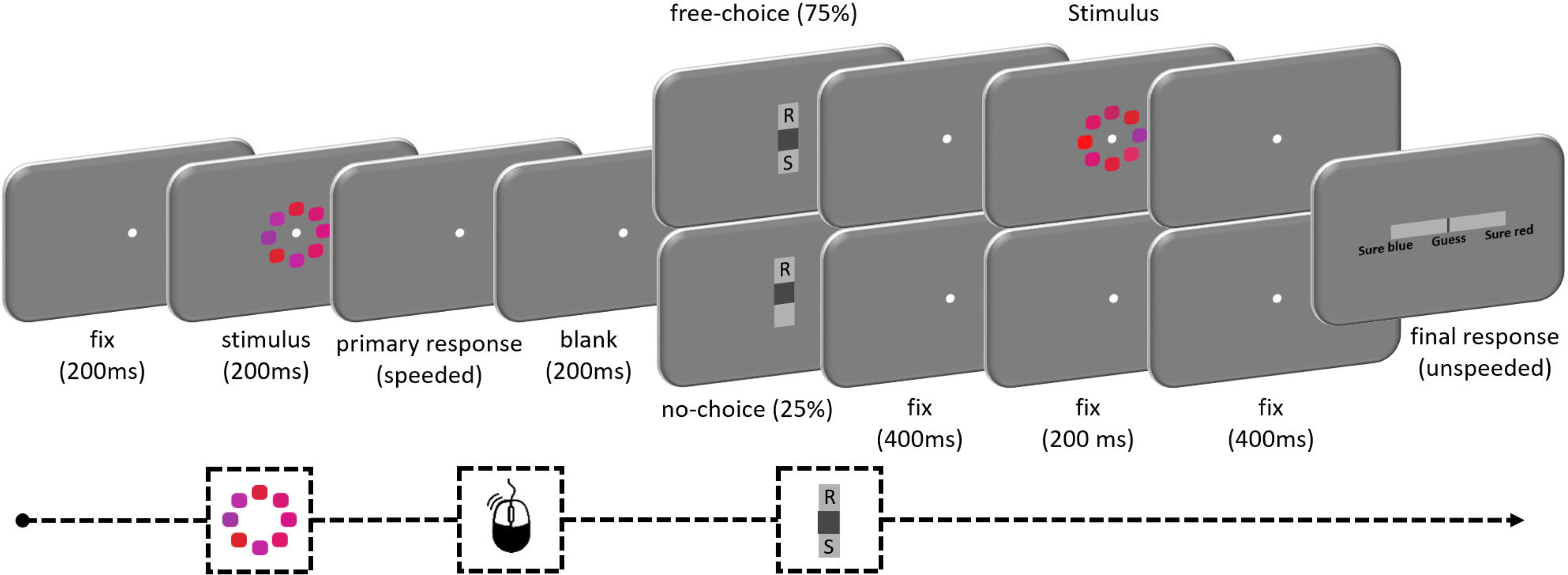
Time line of an experimental trial. A stimulus was presented for 200ms, and participants made a speeded response with the mouse, deciding whether the average color of the eight elements was red or blue. On free-choice trials (75%), participants subsequently used a vertical slider to choose either to see the stimulus again in an easier version (by moving the grey cursor towards S) or to give their response (by moving the grey cursor towards R). When the stimulus is shown again, the mean of the eight elements is more clearly red and the variance is smaller (note that the displayed change is exaggerated for illustration purposes). On no-choice trials (25%), participants could only choose to give their response. Finally, on all trials participants jointly indicated their final response and level of confidence on a horizontal continuous response scale. Being accurate was rewarded (+5 points), errors were punished (−5 points), and there was a small cost associated with sampling more information (−1 point).

When participants chose to see the stimulus again, the values of the stimulus were slightly altered so that the distance to the boundary was higher (*C’* = *C* +/−. 01) and the variability lower (*V’* = *V* –.0167), making the discrimination easier. They were presented with a fixation point for 400ms, this easier stimulus for 200ms and a fixation point for 400ms. When participants opted (or were forced) to give their response without seeing the stimulus again, they simply viewed the fixation point for the same total amount of time (1000ms).

Afterwards, a horizontal response scale appeared (0.4° high and 9.0° wide) with a slider (0.4° high and 0.1° wide) in the center. The left-hand side of the bar was labelled as ‘sure blue’, the right-hand side as ‘sure red’ (depending on the counterbalancing of the response mapping). The location of the slider on the scale was translated into a numerical score, ranging from –50 (sure blue) to +50 (sure red), with every three screen-pixel increments (0.09°) resulting in a difference of one confidence point. Participants moved the cursor with their mouse to indicate jointly their response and their level of confidence, and confirmed by pressing the space bar. A response could not be given when the cursor was exactly in the middle (0 on the scale), so participants were always forced to make the categorical judgment between red or blue. They were instructed to make this judgment at their own pace. Accuracy for the final response was scored as a binary variable (i.e., ignoring confidence level). Confidence was scored as the absolute value on the response scale. To account for between-participant variation in use of the confidence scale and drift in confidence judgments over the course of the experiment, ratings were *z*-scored separately for each participant and each block.

Participants gained 5 points for correct answers and lost 5 points for errors. They could win up to an additional £4.92 by scoring points (650 points = £1). Choosing to see the stimulus again in an easier version cost 1 point, giving participants an incentive to sample more information only when the benefit of doing so (in terms of increasing the probability of making a correct choice) outweighed this cost. Participants were explicitly instructed that they could score more points by strategic use of the see again option.

The main part of the experiment comprised 10 blocks of 64 trials, with balanced numbers of trials for each combination of mean and variance separately for each trial type (free-choice vs. no-choice), in pseudo-randomized order. Each block started with 8 additional practice trials in which the free- vs. no-choice phase was omitted and participants received auditory feedback on the accuracy of their responses. This was done to maintain a stable color criterion over the course of the experiment. Before the main part of the experiment, several practice blocks were administered. In the first block (64 trials), participants practiced the color judgement task, and only were to give a speeded response with the mouse, with auditory feedback to signal decision accuracy. In Blocks 2 and 3 (64 trials each), the second response (including the confidence judgment) was added to the task. No feedback was delivered during these blocks, which served to familiarize participants with the confidence rating scale. Finally, practice block 4 was identical to the main part of the experiment.

At the end of each block (starting from Block 2), the *C* value in the low-mean condition was adjusted depending on performance in that block. Specifically, an inverse efficiency score (median RT/p(correct)) was calculated for the condition with low mean and low variance and the condition with high mean and high variance. When the difference between both was higher or equal than 100/50/10 in absolute value, the *C* value of the low-mean condition was adjusted by .0025/.0012/.0005 respectively, depending on the sign of the difference in order to match performance in these two conditions. This manipulation was carried out to create two conditions with equated performance but different levels of confidence (Boldt, de Gardelle, & Yeung, 2017; Desender et al., 2018). This difference in confidence was not significant in the current data (see Results section), and was not a focus of the current work.

### EEG recording and preprocessing

Participants sat in a dimly lit, electrically shielded room. EEG data were recorded using a fabric cap (QuickCap, Neuroscan) with 32 channels, all referenced to the right mastoid online. Vertical and horizontal electro-oculogram was measured from above and below the left eye and the outer canthi of both eyes. Impedance was kept below 50kΩ. The data were continuously recorded using SynAmps2 amplifiers (Neuroscan), sampled at 1000 Hz. Stimulus-locked data were baselined –100ms to 0ms prior to stimulus onset. In addition, these data were aligned to the time of the primary response, while keeping the same pre-stimulus baseline. Independently from this, the raw data were also locked to the primary response with a baseline –100ms to 0ms prior to response onset. For analyses locked to the information-seeking decision, the response-locked data (keeping the same pre-response baseline) were realigned to the onset of the information-seeking decision (i.e., the space bar press confirming the decision). In preprocessing, segments containing gross artifacts were first identified by visual inspection and removed. Next, eye blinks were removed using Independent Component Analysis (ICA), and segments containing values ±200 mV were excluded using extreme value rejection. Bad (noisy) channels were replaced by an interpolated weighted average from surrounding electrodes using the EEGLAB toolbox (Delorme & Makeig, 2004) in Matlab. Finally, segments containing further artefacts, identified by visual inspection, were removed prior to averaging. For plotting purposes only, data were filtered using a 10 Hz low pass filter.

### Statistical analyses

#### Behavioral analysis

To test whether our different measures of performance lawfully scaled with difficulty, indices of performance were calculated separately for the factors mean and variance. This was done for median RTs on correct trials and mean accuracy of the primary response (calculated based on all trials), mean accuracy of the secondary response, mean confidence on the secondary response (calculated on no-choice data), and the proportion of see again choices (calculated on free-choice data). A repeated measures ANOVA with the factors mean (high or low) and variance (high or low) was then carried out on all these indices.

Next, we fitted a mixed regression model to the data to test whether confidence predicts information seeking over and above other factors. This analysis cannot be carried out at the trial-level, because confidence and see again choices are measured on separate parts of the data (no-choice versus free-choice, respectively). Therefore, for all variables we computed eight data points (2 levels of mean × 2 levels of variance × 2 colors), separately for each participant. Specifically, we computed (i) mean confidence based on the no-choice data, (ii) the proportion of see again choices based on the free-choice data, (iii) mean accuracy of the primary response based on all data, and (iv) median RTs on the primary response based on all data, separately for the factors evidence variance (high or low), evidence mean (high or low), and color (red or blue). Values were calculated separately for each color to partition the data in a more fine-grained manner, but this variable was not taken into account in the analyses. For ease of interpretation, low variability and high mean were dummy coded as reference categories so that a positive effect of each factor corresponds to an increase in difficulty. We used mixed regression modelling (using the lme4 package in R; Bates, Maechler, Bolker, & Walker, 2015) to construct models of increasing complexity. For each model, random slopes were added for all variables for which this significantly increased the fit compared to that model without random slopes. When required, degrees of freedom were estimated using Satterthwaite’s approximation (using the lmerTest package; Kuznetsova, Brockhoff, & Christensen, 2014).

#### Event-related potentials

In a first set of analyses, we wanted to confirm that our data showed the usual modulation of pre- and post-decisional event-related components as a function of decision confidence. Building on previous work, event-related potential (ERP) indices of confidence and see again choices were tested at electrode CPz (Boldt & Yeung, 2015). Significant time windows during which ERPs for high and low confidence (or for respond and see again trials) differed from each other were identified using a standard two-tailed within-subjects cluster-based permutation test using custom code in Matlab. Elements that were adjacent and significant (element-level *p* < .05) were collected in a cluster. Cluster-level test statistics consisted of the absolute sum of *t*-values within each cluster, and these were compared with a null distribution of test statistics created by drawing 1000 random permutations of the observed data. A cluster was considered significant when its (cluster-level) *p*-value was below .05. To examine whether the modulation of the ERPs by confidence extended over and above our manipulation of task difficulty, single-trial mean amplitudes within the significant time windows were extracted and submitted to separate linear mixed regression models. Models of different complexity were estimated using the factors variance and mean. Confidence was then added to the winning model, testing whether it predicts EEG activity over and above task difficulty.

#### Time-resolved decoding

To test for overlap in the neural coding of confidence and see again choices, a classifier was trained separately for each participant using single-trial logistic regression based on the linear derivation method introduced by Parra et al. (2005). This approach identifies the spatial distribution of scalp EEG activity in a given time window that maximally distinguishes two classes to deliver a scalar estimate of component amplitude. Two sets of analyses were performed. First, it was tested how well information-seeking choices can be decoded from EEG data (within-condition decoding). This analysis was a first step in identifying the time window during which information-seeking choices are decodable. Both the training and testing set were restricted to trials with correct responses on the primary decision only, in order to dissociate between the coding of information seeking and coding for errors. A 10-fold cross validation approach was used to avoid overfitting. To increase reliability, this 10-fold cross validation approach was repeated 100 times, resulting in 1000 classification scores per time point. These scores were then averaged, and below we report these averaged classification values. For the second set of analyses, we tested how well see again choices can be decoded from a classifier trained to discriminate EEG activity associated with differing levels of confidence. Confidence judgments on no-choice trials were median split into high and low values. The decoder was then trained to discriminate high vs. low confidence judgments (on correct trials only, from no-choice trials) and tested on see again choices on correct trials from free-choice data; thus there was no overlap between training and testing data. This approach was repeated 1000 times. Variability across iterations arose due to the random selection of data necessary to obtain an equal number of high and low confidence trials in the training data and an equal number of see again and respond trials in the free-choice data.

To increase the signal-to-noise ratio of data used for decoding, classifiers were trained on time-averaged signals within discrete temporal windows (window width of 106ms, moving in 10ms increments along entire epochs aligned to stimulus onset, initial decisions and see again choices). The ability to successfully classify individual trials was quantified by calculating the Az score, which gives the area under the receiver operating characteristic (ROC) curve, derived from signal detection theory (Stanislaw & Todorov, 1999). Classifiers were trained and tested on each time point of the EEG data. Thus, this method produces a 2D (training time × testing time) decoding performance matrix. Specific dynamics of mental representations can be unraveled by evaluating the shape of the decoding matrix (King & Dehaene, 2014). The statistical reliability of this decoding was determined via a bootstrap procedure, comparing actual classification with classifier performance on trials with randomized condition labels (1000 iterations with different randomizations), to provide an estimate of the null classification. Clusters were formed via paired-samples *t-*tests for the entire 2D matrix, comparing true and null classifications. Neighboring elements that passed a threshold value corresponding to an (element-level) *p* value of 0.01 (two-tailed) were collected into a separate cluster. Elements were considered as neighbors when they were (cardinally or diagonally) adjacent. Cluster-level test statistics consisted of the absolute sum of t values within each cluster, and these were compared with a null distribution of test statistics created by drawing 1000 random permutations of the observed data. A cluster was considered significant when its (cluster-level) *p*-value was < .05.

## Results

### Behavioral results

#### Performance, confidence and information seeking

We first confirmed that all behavioral measures (performance, confidence and information seeking) scaled with task difficulty (see Figure 2). For the primary perceptual decision (speeded color discrimination) data, a repeated measures ANOVA showed that both median RTs on correct trials and mean accuracy were significantly affected by color mean (RTs: *F*(1,14) = 14.96, *p* = .002; accuracy: *F*(1,14) = 72.94, *p* < .001), and by color variance (RTs: *F*(1,14) = 27.89, *p* < .001; accuracy: *F*(1,14) = 38.37, *p* < .001), but not by their interaction (RTs: *F* < 1; accuracy: *F* < 1). Mean accuracy of the final response (irrespective of the level of confidence), calculated on the data of the no-choice condition in which participants were forced to respond without viewing the stimulus again, was likewise affected by both color mean, *F*(1,14) = 45.55, *p* < .001, and color variance, *F*(1,14) = 23.56, *p* < .001, but the interaction failed to reach significance, *F*(1,14) = 4.32, *p* = .056. Mean confidence, again calculated on the data of the no-choice condition (to avoid effects of seeing the stimulus again), was affected by color mean, *F*(1,14) = 32.18, *p* < .001, and color variance, *F*(1,14) = 21.73, *p* < .001, but not by their interaction, *F* < 1. Finally, on free-choice trials, participants on average chose to see the stimulus again in an easier version on 43.2% (range 2% – 94%) of the trials. The proportion of see again choices was affected by mean, *F*(1,14) = 21.75, *p* < .001, and variance, *F*(1,14) = 20.02, *p* < .001, with no significant interaction between these factors, *F*(1,14) = 3.22, *p* = .09. The large variation in see again choices did not correlate with individual differences in overall mean confidence on forced choice trials, *r*(13) = .22, *p* = .428, *BF* = .26, or accuracy on the primary response, *r*(13) = .36, *p* = .18, *BF* = .47. Replicating previous work (Desender et al., 2018), participants choose more often to see the stimulus again in the high mean – high variance compared to the low mean – low variance condition, *t*(14) = 2.47, *p* = .027, despite similar primary accuracies in both conditions, *p* = .189, but note that RTs were also slower in the former, *t*(14) = –2.819, *p* = .014. Unexpectedly, there was no difference in confidence between these conditions, *p* = .570, perhaps reflecting the slight speed-accuracy tradeoff difference between these conditions, with slightly more cautious responding in the high mean –high variance condition.

**Figure 2.**
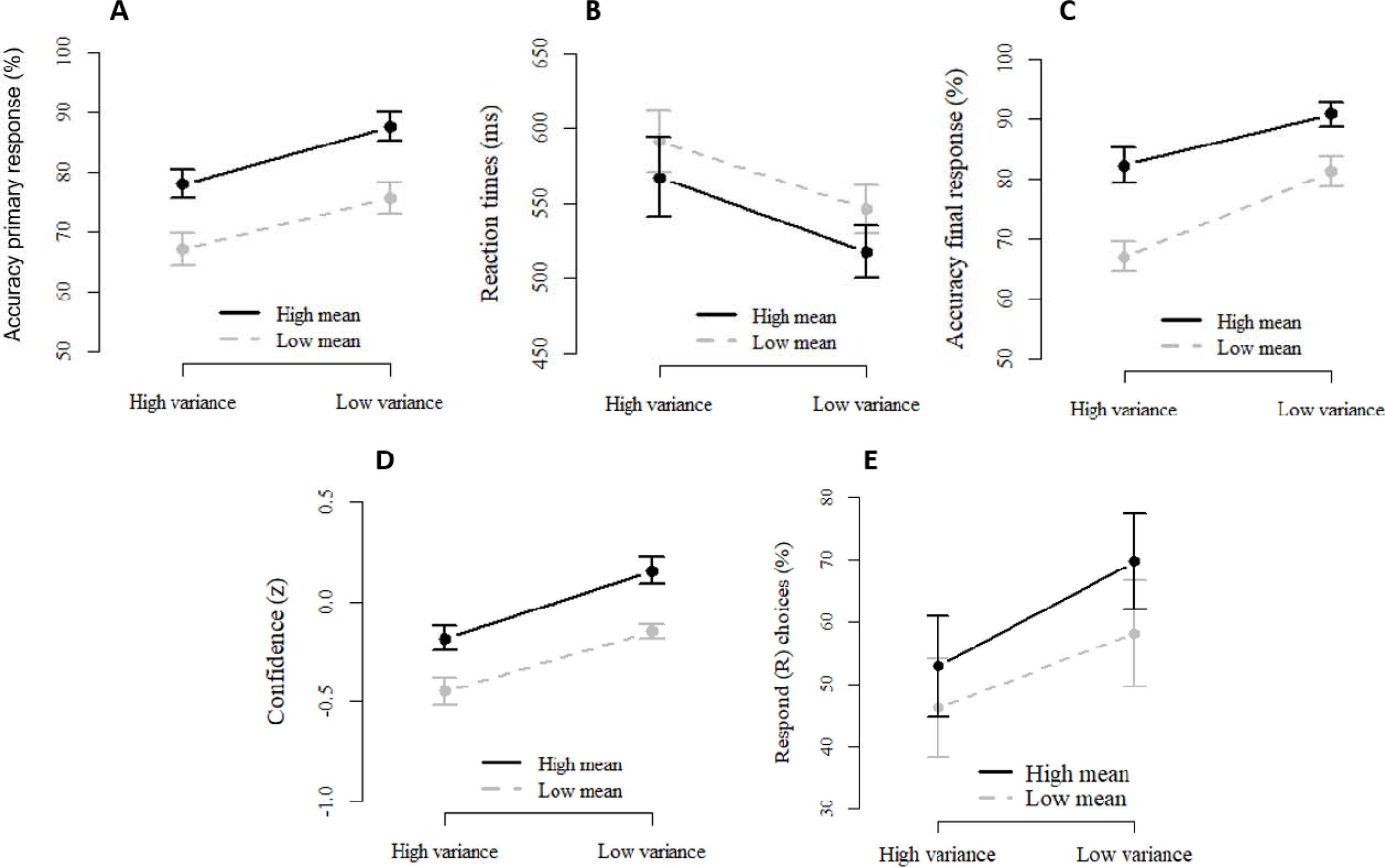
Behavioral performance is modulated by mean and variance. A. Mean accuracy of the primary response (based on all data). B. Median reaction times of the primary response (based on all data). C. Mean accuracy of the final response (based on no-choice data). D. Mean standardized confidence (based on no-choice data). E. The number of trials on which participants waived the see-again option (based on free-choice data).

#### Behavioral confidence predicts information seeking

To further interrogate potential sources of variability in observed information seeking behavior, mixed regression models of increasing complexity were fit to the data predicting variation in the proportion of see again choices across conditions of the experimental design (see Methods) by five predictors: the factors color variance (high or low) and color mean (high or low), and the variables median primary RTs (both on free-choice and no-choice), mean accuracy of the primary response (both on free-choice and no-choice) and mean confidence (on no-choice trials). A model building strategy was used (see Table 1). The experimental variables mean and variance explained a significant part of the variance in information seeking (Model 1), and measures of primary task performance (accuracy and RT) significantly increased the fit (Model 2). Crucially, adding confidence to a model that already contained these four variables provided the best fit to the data. As predicted, in this final model (Model 3), there was a clear negative effect of confidence on see again choices, *β* = –.08, *t*(85.63) = –3.09, *p* = .003, whereas the effects of RT, *β* = .15, *p* = .065, and variance, *β* = .03, *p* = .069, were no longer statistically significant. The effects of accuracy and mean were not significant either, both *p*s > .10. Thus, although mean, variance, RTs and accuracy explained a significant amount of variation in information seeking in simpler models, their statistical contributions were accounted for by confidence.

**Table 1.**
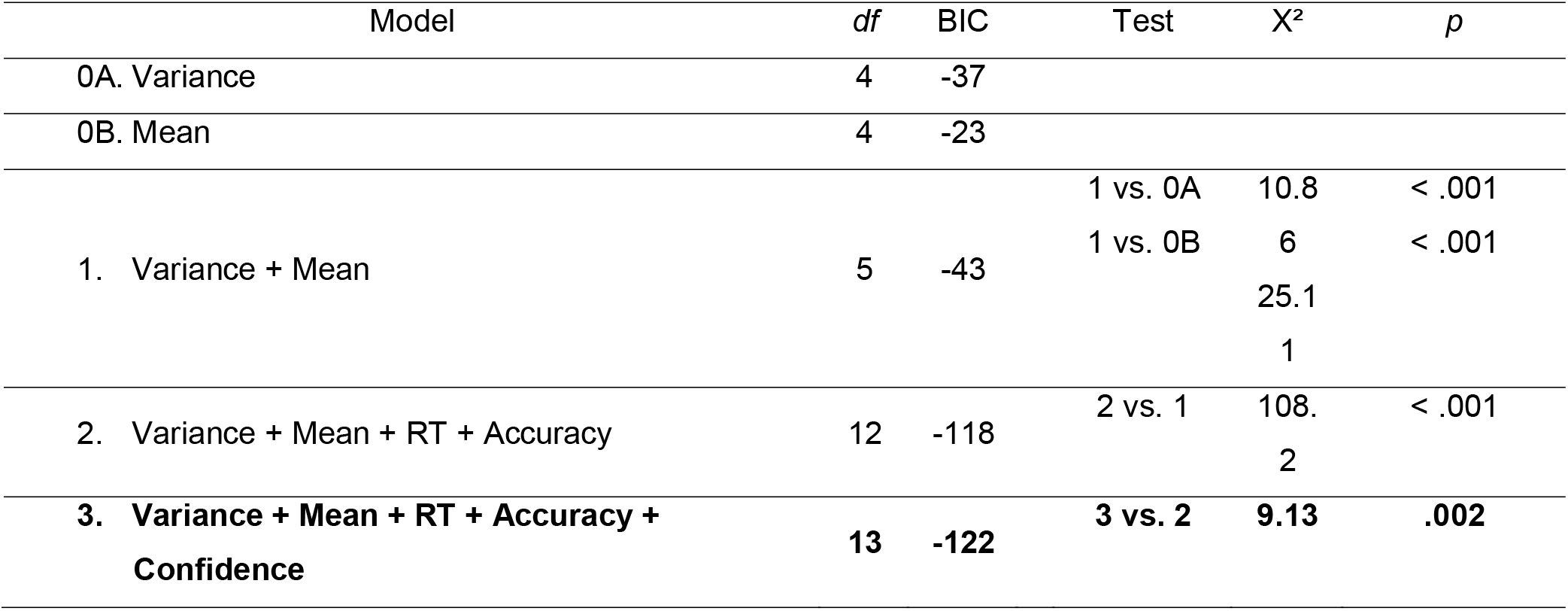
Models of different complexity predicting information seeking. Winning model is indicated in bold face.

### EEG analysis

#### ERP markers of confidence

To examine how ERP waveforms are modulated by confidence, correct trials in the no-choice condition were split into high and low confidence bins via median-split separately for each participant. As can be seen in Figure 3A, at electrode CPz the stimulus-locked ERP showed significant modulation by confidence from 414ms to 581ms, *p* = .016 (cluster level), corresponding to a typical P300 component (Hillyard, Squires, Bauer, & Lindsay, 1971; Polich, 2007). Mean amplitude was more positive for trials on which participants later on indicated high (vs. low) confidence. By contrast, the ERP aligned to and following the primary response showed the opposite pattern: more negative amplitudes for high confidence trials were observed from 403ms post-response until the end of the analyzed epoch (700ms; *p* = .008, cluster level), corresponding to a typical Pe component (Ridderinkhof, Ramautar, & Wijnen, 2009). The topographies associated with these significant time windows showed that both effects had a similar centro-parietal scalp distribution.

**Figure 3.**
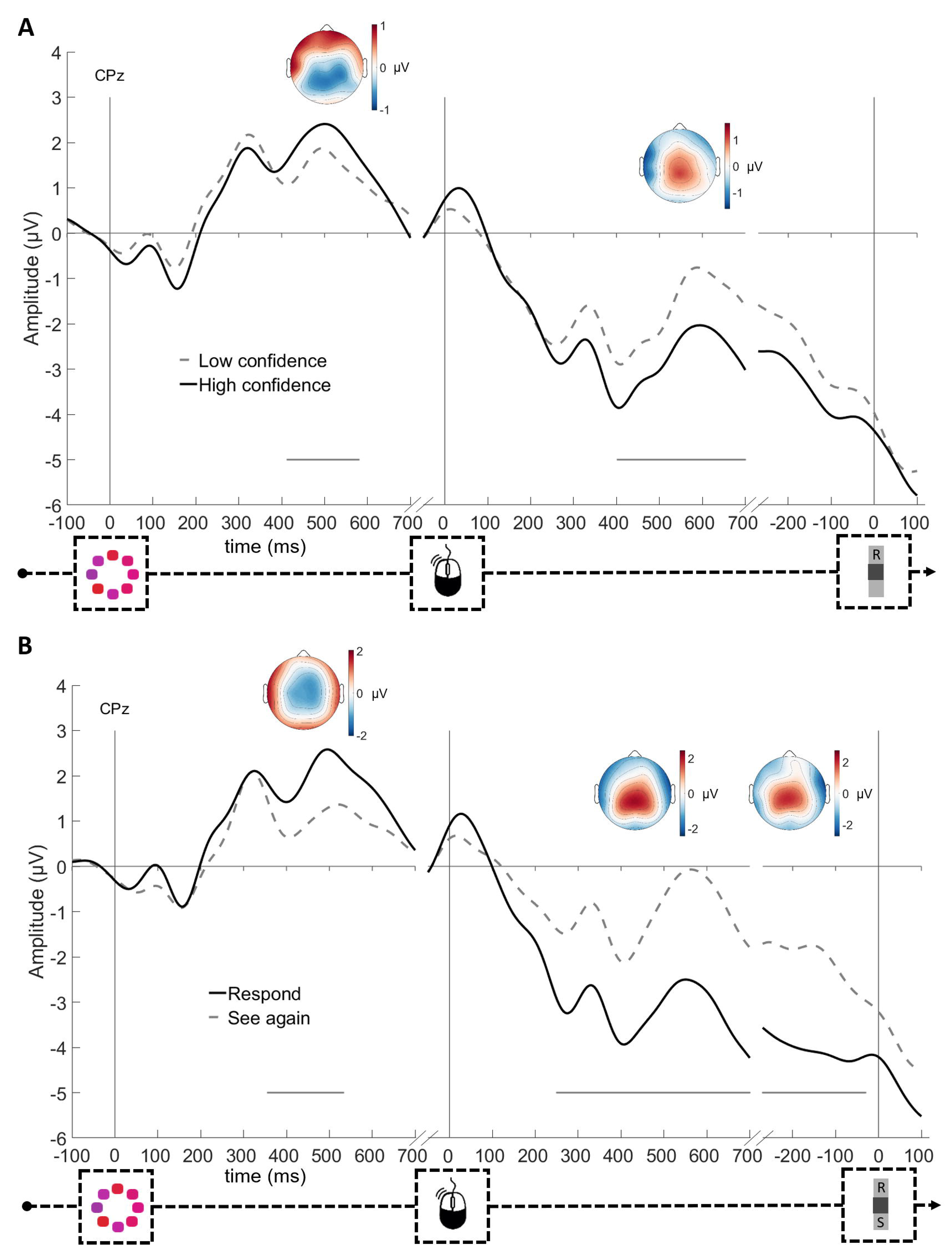
Event-related potentials as a function of confidence (A) and information-seeking choices (B). Head plots show the difference in scalp distribution during the significant time periods for low-high confidence (A) and see again – respond (B). Grey horizontal lines reflect clusters during which both conditions significantly differ. Note that high and low confidence are calculated from no-choice data and information-seeking choices from free-choice trials.

#### Post-decisional ERPs are modulated by confidence over and above task condition

We next examined whether the modulation of the ERPs by confidence extended over and above our manipulation of task difficulty. To do so, single-trial mean amplitudes within the significant time windows were extracted and submitted to separate linear mixed regression models. Models of different complexity were estimated, in which mean amplitude was predicted by the factors color variance, color mean, and their interaction, and it was then tested whether confidence significantly improved the fit of the winning model (as it did in the analysis of information seeking behavior described above).

First, we examined the significant modulation in the stimulus-locked ERPs, from 414ms to 581ms (i.e., the P3). As can be seen in Table 2, a model with only variance provided a better fit to the data (lower BIC value) than any of the models including mean. Crucially, adding confidence as a predictor to this null model did not improve the fit (Model 3). This analysis suggests that the modulation of the P3 by confidence is driven by differences in signal quality, as induced by the variance of the presented stimulus elements.

**Table 2.** Model comparisons for the stimulus-locked cluster modulated by confidence, from 414ms until 581ms stimulus-locked, reflecting the P3 component. Winning model is indicated in bold face.

| Model | df | BIC | Test | X<sup>2</sup> | p |
| --- | --- | --- | --- | --- | --- |
| **0A. Variance** | **4** | **11258** |  |  |  |
| 0B. Mean | 4 | 11268 |  |  |  |
| 1. Variance + Mean | 5 | 11264 | 1 vs.<br>0A | 2.09 | .15 |
| 2. Variance * Mean | 6 | 11270 | 2 vs.<br>0A | 3.17 | .20 |
| 3. Variance + Confidence | 5 | 11264 | 3 vs.<br>0A | 2.45 | .12 |

Second, we examined the significant modulation in the response-locked ERPs, from 403ms until the end of the epoch (700ms) post-response (i.e., the Pe). The model comparison reported in Table 3 indicates that a model featuring variance provides a better fit than a model featuring mean (lower BIC value), and that neither model including mean increases the fit. Crucially, adding confidence to the null model with variance significantly increased the fit (Model 3). In the winning model, both confidence, *F*(1,1699.7) = 7.96, *p* = .005, and variance, *F*(1,1697.7) = 22.51, *p* < .001, explained significant variation in Pe amplitude. Thus, this analysis demonstrates that, in contrast to the stimulus-locked P3 cluster, modulation of the response-locked Pe cluster by confidence went beyond the effect of experimentally-induced task difficulty.

**Table 3.** Model comparisons for the response-locked cluster modulated by confidence, from 403ms until 700ms post-response, reflecting the Pe component. Winning model is indicated in bold face.

| Model | df | BIC | Test | X<sup>2</sup> | p |
| --- | --- | --- | --- | --- | --- |
| 0A. Variance | 4 | 10887 |  |  |  |
| 0B. Mean | 4 | 10911 |  |  |  |
| 1. Variance + Mean | 5 | 10892 | 1 vs.<br>0A | 3.11 | .078 |
| 2. Variance * Mean | 6 | 10899 | 2 vs.<br>0A | 3.15 | .21 |
| **3. Variance + Confidence** | **5** | **10885** | **3 vs.**<br>**0A** | **7.97** | **.005** |

#### ERP markers of information-seeking choices

Figure 3b shows the ERP waveforms at CPz, separately for see again and respond directly choices, again averaged over correct trials only. The results closely mirror the modulation of the ERPs by confidence. The stimulus-locked ERPs showed more positive amplitudes on respond compared to see again trials, from 357ms until 534ms, *p* = .028 (cluster level). The response-locked ERPs showed more negative amplitudes on respond compared to see again trials, from 250ms until the end of the epoch (700ms, *p* < .001, cluster level). Finally, the latter difference remained significant up until 29ms before the information seeking decision, *p* = .008 (cluster level). Similar to confidence, all significant effects had a clear centro-parietal scalp distribution, although the stimulus-locked P3 component for information-seeking choices has a slightly more anterior scalp distribution than that for confidence.

In sum, analysis of the ERPs provides preliminary evidence for a link between confidence and information seeking, given that both processes have very similar neural markers. In the following, we use multivariate single-trial decoding to provide a more rigorous appraisal of this link, testing the informational content of these neural markers (i.e., whether information-seeking choices can be decoded reliably) and whether neural markers of confidence are predictive of information-seeking behavior.

### Time-resolved decoding

#### Within-condition decoding of information-seeking choices

We first tested whether information-seeking choices can be decoded over time from the EEG data. This analysis is a first step in identifying the time window during which information-seeking can be decoded. In this analysis, classifiers were both trained and tested on data from free-choice trials, using 10-fold cross-validation as described above. Decoders were trained and tested on each point in time, thus shedding light on the generalization of the discriminate pattern over time. In the stimulus-locked matrix (Figure 4A), no robust decoding was possible in the pre-stimulus period, all cluster *p*s > .143. Note that the decoder was only trained on data up until 400ms post-stimulus (13^th^ percentile of RTs) to avoid contamination from post-response data (from trials with short RTs). To complement this, the same analysis was repeated after the data were realigned to the time of the response (but keeping the same pre-stimulus baseline). Again, no decoding was visible pre-response, all *p*s > .156, but there was a significant cluster that largely fell post-response *p* = .040 (Figure 4B). Next, we decoded EEG data measured in the post-response period, using a conventional pre-response baseline. This analysis revealed that, in marked contrast to the pre-response epoch, it was possible to classify see again choices reliably across the entire epoch from the initial task response to the subsequent see again choice: There was a highly significant cluster that began just prior to primary response execution (Figure 4C; *p* < .001, cluster level), and peaked just prior to the decision of whether or not to sample more information (Figure 4D; *p* < .001, cluster level). Decoding performance was strongest along the diagonal, indicating best decoding when the cross-validation test data came from the same time period as the classifier training data. Nevertheless, both clusters also displayed significant off-diagonal decoding, indicating that the cross-validation test data could be predicted above chance level even when classifier training data was obtained from a different time window. This strong temporal generalization suggests a neural activity pattern that is consistent and sustained over time during the formation of information-seeking choices.

In sum, information-seeking choices could be decoded from EEG data, but only during the time window following the primary task response. No above-chance decoding was visible when classifiers were trained on EEG data from the pre-response window.

**Figure 4.**
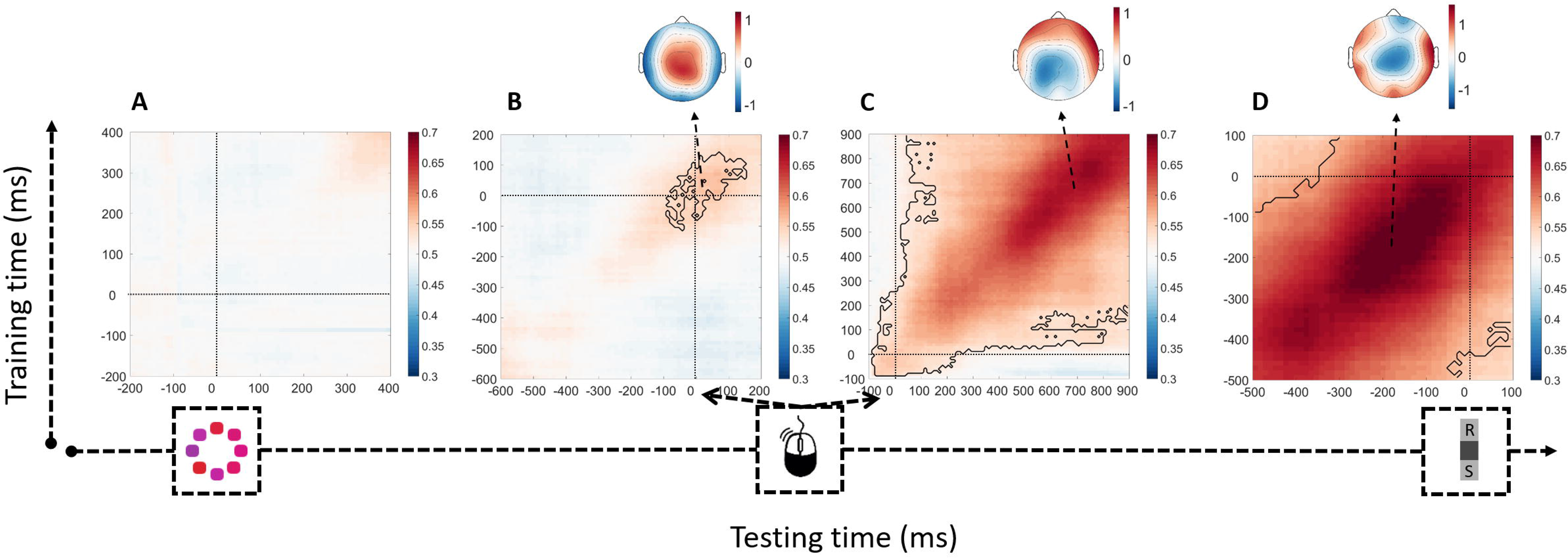
Within-condition decoding of information-seeking choices. Classifiers were trained and tested on all time points (on correct trials only; steps of 10ms and a sliding window of 106ms). The topographies display the scalp projections obtained from the logistic regression classifier at the training time where classification is maximal. Panels A and B have a pre-stimulus baseline (−100ms until 0ms) and panels C and D have a pre-response baseline (−100ms until 0ms). Solid black lines indicate significant clusters (p < .05). Note that the training times of each panel correspond to the testing time of that panel; for example, t = 0 corresponds to stimulus, response, response, and information-seeking decisions in panels A-D, respectively.

#### Across-condition decoding of information-seeking choices by confidence

While the previous analyses already hinted at the importance of post-decisional neural activity for information-seeking choices, those analyses are uninformative about the role of confidence in this process. Our final set of analyses provided the critical, direct test of the hypothesis that confidence underpins adaptive information seeking. Specifically, we tested whether an EEG classifier trained to decode confidence on no-choice trials (i.e., in which participants were not asked to decide whether or not to seek further information before their final decision) would be able to predict see again choices on free-choice trials (i.e., on the separate set of trials in which participants had the option of seeking or declining additional information). Importantly, on no-choice trials participants did not make an information sampling choice themselves, but were forced to select the option to give their response. This rules out the possibility that our classifier is decoding incidental processes related to this choice, such as motor-related neural activity resulting from the information-seeking choice. Moreover, to ensure that our classifier was decoding decision confidence, and not achieving above-chance classification by exploiting other correlated features of the data (such as trials in which participants changed their mind), only trials on which participants were correct in both their initial and their final decision were used to train the classifier.

In the resulting stimulus-locked matrix, there was no sign of above-chance decoding, all *p*s > .289 (see Figure 5A). A complementary analysis in which these data were realigned to the time of the response (keeping the same pre-stimulus baseline) also showed no reliable pre-response decoding, all *p*s > .218 (Figure 5B). The response-locked data, by contrast, showed significant decoding from around 350 ms to 700 ms after the response, *p* = .008 (cluster level; Figure 5C). Thus, a decoder trained to predict confidence from EEG data from no-choice trials was able to predict information seeking decisions from EEG data recorded on free-choice trials, specifically in this post-response window. As becomes clear from the associated topography in Figure 5C, the decoder largely relied on centro-parietal electrodes to make its predictions. Significant decoding was not observed in analysis of data time-locked to the information sampling decision (Figure 5D, all cluster *p*s > .256).

**Figure 5.**
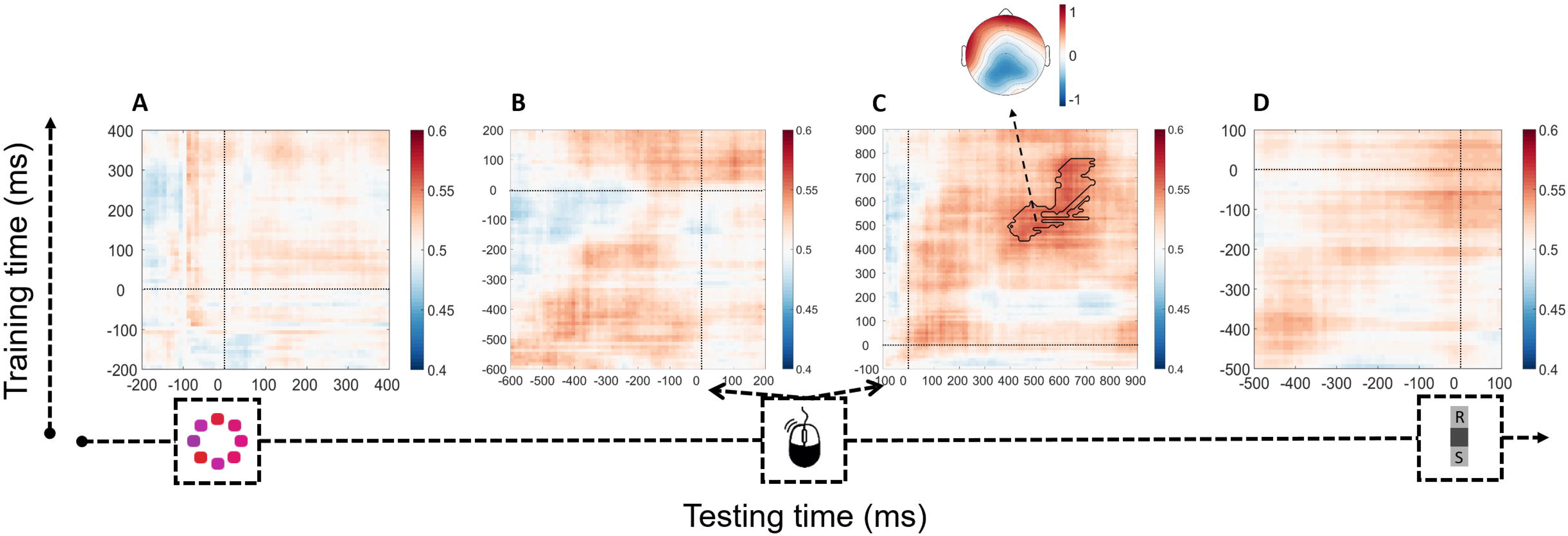
Across-condition decoding of information-seeking choices by confidence. Classifiers are trained on high vs. low confidence from no-choice data and tested on see again vs. respond decisions from free-choice data (both on correct trials only). Above-chance decoding only occurs post-response. The same conventions as in Figure 4 apply.

## Discussion

The current study examined whether neural markers of decision confidence are predictive of information seeking behavior. To this end, we recorded scalp EEG while participants performed a task in which they first made an initial decision about a stimulus, then chose whether or not to sample more information, before providing their final response and level of confidence. Information-seeking choices and confidence similarly modulated pre-decisional (P3) and post-decisional (Pe) ERP components. Using multivariate classification, we then showed that information-seeking choices could be decoded from EEG data from the time of the initial decision to the time of the subsequent information-seeking choice (within-condition decoding). However, no above-chance decoding was visible preceding the initial decision. Crucially, a classifier trained to decode high versus low confidence generalized to prediction of information-seeking choices (across-condition decoding), and this too was restricted to a post-response time window. The time period during which we observed robust across-condition generalisation (i.e., from no-choice trial confidence ratings to free-choice trial information-seeking behavior) corresponds to that of a post-decisional neural marker of decision confidence, suggesting the latter reflects a neural process integral to translating one’s subjective sense of confidence into overt decisions to sample more information.

When confronted with difficult decisions, humans seek further information to improve the quality of their decisions. Unsolicited information does not affect decisions, whereas decisions are more accurate when additional information is actively solicited (Yoong & Hung, 2010). This suggests that the act of seeking further information is driven by an internal evaluation. A straightforward hypothesis is that humans internally compute the probability of making a correct decision, and when this probability is low they seek additional information. Indeed, in a recent behavioral study we were able to demonstrate that explicitly represented decision confidence predicts information seeking, even across conditions matched for objective difficulty (Desender et al., 2018). Consistent with these earlier findings, here we found that decision confidence, not objective accuracy or stimulus difficulty, was the main variable predicting information seeking in a mixed model analysis. The current work significantly extends these previous behavioural findings by characterising the neural signatures integral to translating decision confidence into overt information-seeking choices. In particular, we could predict information seeking behavior based on confidence; however, we could do so only in post-decisional EEG activity, even though averaged ERP waveforms varied significantly as a function of confidence and see again choices also in the pre-decisional period. As such, our findings converge with theoretical work arguing that post-decisional evidence accumulation plays a critical role in confidence judgments (Fleming & Daw, 2016; Moran, Teodorescu, & Usher, 2015a; Pleskac & Busemeyer, 2010) and subsequent actions: Depending on the strength of the post-decisional evidence (i.e., reflecting the degree of confidence), participants will seek additional information or not.

The post-decisional neural marker observed in the current work closely resembles the classical Pe component of the ERP (Ridderinkhof et al., 2009). This neural marker has been suggested to reflect an evidence accumulation signal evaluating the likelihood that the just-executed response was incorrect (Murphy et al., 2015; Steinhauser & Yeung, 2010). Accordingly, the amplitude of this signal (reflecting the amount of evidence for an error) has been shown to scale inversely with decision confidence (Boldt & Yeung, 2015). Our observation of this signal after participants made a primary decision in both the no-choice and free-choice conditions suggests that, in both conditions, evidence is accumulated about the likelihood of this decision being correct. This evaluation process could in principle be used to guide both immediate, binary information-seeking choices (by imposing a single threshold on the evolving tally of error evidence; Murphy et al., 2015) and later confidence reports (by translating the final continuous evidence tally into a correspondingly graded expression of confidence). The present finding of significant across-condition decoding provides important support for this idea by showing that both confidence reports and information-seeking choices appear to be governed by the same underlying neural signal.

An alternative interpretation of our results could be that the post-decisional neural marker we describe reflects the internal decision to seek more information, rather than a common signal that can be leveraged for making both information-seeking decisions *and* graded confidence reports. In other words, perhaps participants made internal information-seeking decisions on no-choice trials despite there being no explicit requirement to do so, and the neural signal that is central to our analyses reflects this. Such an explanation of our findings seems unlikely for two reasons. First, our post-decisional neural signal highly resembles the well-characterized Pe component, both in time and scalp distribution (Ridderinkhof et al., 2009). The Pe is reliably observed in paradigms that do not require any overt post-decisional response, which makes it unlikely that it reflects a signal specifically related to this additional response. Second, if this post-decisional neural signal directly reflects the information sampling choice, it should be related to systematic biases that are observed in such choices. We indeed observed large inter-individual differences in the tendency to prefer one of the information-seeking options, with some participants biased towards respond choices (*N* = 10, on average 23% see again choices, RT_see again_ = 1245ms vs. RT_respond_ = 800ms, *t*(9) = 4.18, *p* = .002) and others towards see again choices (*N* = 5, on average 73% see again choices, RT_see again_ = 797ms vs. RT_respond_ = 953ms, *t*(4) = –2.02, *p* = .11). For both subgroups, however, the post-decisional neural marker was of higher amplitude when participants chose to see more information (replicating Figure 3B; both *p*s < .001). This observation is hard to reconcile with the idea that post-decisional neural activity reflects processes related specifically to the information-sampling choice, but are compatible with our interpretation that this activity indexes an evaluation of the accuracy of the just-executed response, which then informs explicit confidence reports and information-seeking choices.

Neither within-condition nor across-condition decoding analyses yielded above-chance decoding of information-seeking choices in the time window occurring before the response. This lack of decoding is striking given that pre-decisional neural activity (corresponding to the well-studied P3 component) was found to be sensitive to confidence and information-seeking in univariate analyses. In this regard, it is important to note that not all neural signals that are modulated by confidence are also involved in the representation of explicit (i.e., subjective) confidence (Pouget, Drugowitsch, & Kepecs, 2016). Several studies showed predecisional neural markers that are sensitive to the level of confidence (Gherman & Philiastides, 2015, 2017; Kiani & Shadlen, 2009; Odegaard et al., 2018). For example, Odegaard and colleagues (2018) demonstrated that opt-out decisions (presumably reflecting low confidence) are associated with weak traces of neural evidence in monkey superior colliculus occurring before the response. However, using a positive-evidence manipulation that dissociates evidence quality from confidence, they were able to show that the pre-decisional neural activity tracks evidence quality, not subjective confidence (both of which are typically closely associated). A similar observation was made in the current work: Significant modulation of the pre-decisional neural marker by confidence disappeared once stimulus variability (i.e., a measure of evidence quality) was taken into account. Together, this might explain why pre-decisional neural markers are modulated by confidence, but do not predict information-seeking choices.

Our findings are of relevance to research on the role of uncertainty in action control (Daw, Niv, & Dayan, 2005; Yu & Dayan, 2005). From a Bayesian perspective, uncertainty (i.e., the inverse of confidence) can be used as a cue for behavioural control (Daw et al., 2005). From this, it follows that decision confidence should predict strategic decisions, such as whether or not to sample more evidence (Meyniel et al., 2015). Our findings are the first empirical demonstration that decision confidence and the adaptive act of information-seeking share the same neural signal. Another relevant connection to our work is research on exploitation-exploration dilemmas. When forced to decide from which of several patches to harvest, participants are faced with the dilemma between exploiting a known patch or exploring unknown but potentially more rewarding patches. Empirical and modelling work has demonstrated that participants’ uncertainty about these choices arbitrates between both strategies. Consistent with normative theories (Sutton & Barto, 2018), participants actively explore unknown patches to gain information and reduce their uncertainty (Badre, Doll, Long, & Frank, 2012; Cavanagh, Figueroa, Cohen, & Frank, 2012). Interestingly, recent research has indicated an important role for subjective confidence in this process (Boldt, Blundell, & De Martino, 2017). Specifically, when confidence in value representations was low, participants actively explored unknown patches. Although the underlying construction of confidence may differ (confidence in value representation vs. confidence in the accuracy of a decision), in both cases low confidence triggers the need for information-seeking.

## Conclusion

It was shown that a classifier trained to decode confidence based on EEG data was able to predict information-seeking decisions, specifically in the time window following a speeded decision about the stimulus. This suggests that post-decisional neural processes are integral to translating decision confidence into overt decisions to seek further information.

## Funding

K.D. is an FWO [PEGASUS]² Marie Skłodowska-Curie fellow (grant number 12T9717N), and was supported by grants of the Research Foundation Flanders, Belgium (FWO-Vlaanderen; grant numbers 11H3415N and V447115N). A.B. was supported by an Economic and Social Research Council UK PhD studentship.

## Acknowledgments

Thanks to Kerstin Fröber for assistance with EEG prepping, and to Mehdi Senoussi for advice on the decoding analyses.

